# Thalamic nuclei insights into Alzheimer’s disease

**DOI:** 10.64898/2026.05.26.728015

**Authors:** Julie P. Vidal, Daniel J Myall, Jérémie Pariente, Toni L Pitcher, Reece P Roberts, Erin E Cawston, Campbell Le Heron, Tim J Anderson, Catherine A Morgan, Tracy R Melzer, Ian J Kirk, Lynette J Tippett, Patrice Péran, John C Dalrymple-Alford

## Abstract

**INTRODUCTION:** Thalamic nuclei support multiple cognitive processes, yet their integrity in biologically-defined Alzheimer’s disease (AD) remains unknown.

**METHOD:** Amyloid status was determined using PET Centiloids >24 in 1,327 participants from ADNI. Combined with clinical diagnosis, this yielded six groups: amyloid-negative or positive CN-MCI-dementia/AD. Thalamic nuclei volumes were extracted from T1-weighted MRI using the HIPS-THOMAS algorithm.

**RESULTS:** Large volume reductions in the anteroventral, mediodorsal, and pulvinar nuclei were observed in amyloid-positive MCI and AD. Reduced volumes were also evident in amyloid-positive CN, supporting preclinical AD. Adding the anteroventral nucleus improved cognitive status classification in Random Forest analyses. A phenotypic model integrating thalamic nuclei clearly distinguished amyloid-positive groups from amyloid-negative CN and reclassified non-AD patients with 68% of amyloid-negative MCI subjects as CN-like, and 27% of amyloid-positive CN as MCI-like.

**DISCUSSION:** Thalamic volumetry from conventional T1-weighted MRI enhances clinical insight into AD and provides a practical biomarker for disease intervention.

## 1. Introduction

Thalamic nuclei are critical hubs that dynamically regulate multiple neural networks to support cognitive, sensory, motor, and affective functions [1,2]. In Alzheimer’s disease (AD), post-mortem studies suggest that amyloid plaques and hyperphosphorylated tau in the anterior, laterodorsal and intralaminar thalamic nuclei may predate extensive involvement of the hippocampal system [3-7]. AD neuropathology is also found in the mediodorsal and ventral thalamic nuclei, though this is generally less severe [3,5].

Typically, in vivo MRI studies treat the thalamus as a single structure due to poor intra-thalamic contrast [2]. The advent of a FreeSurfer probabilistic atlas of thalamic subregions combined with T1-weighted (T1w) imaging has revealed reduced volume of thalamic nuclei in AD, although the specific pattern of these changes has been inconsistent across studies [8-13]. Subsequent improvements in thalamic nuclei volume delineation were achieved with the introduction of the THOMAS algorithm, which is based on the Krauth-Morel atlas [14] and uses white matter-nulled images to segment ten thalamic nuclei, the habenula, and the mammillothalamic tract [15-17]. The recent addition of histogram-based polynomial synthesis (HIPS-THOMAS) to simulate high-contrast white matter-nulled images from conventional T1w MRI has further enhanced the delineation of thalamic nuclei [18-21]. Critically, this HIPS-THOMAS method provides the scalability necessary for assessing large repositories of MRI data. Initial work using HIPS-THOMAS reports that atrophy of the anteroventral, pulvinar, centromedian, and mediodorsal thalamic nuclei is evident in clinical MCI, expanding to other nuclei with the progression to dementia [22,23]. Despite these advances, studies have often relied on small to medium sample sizes, clinical diagnostic criteria and, bar one study, have lacked biomarker confirmation. That study examined 14 MCI patients with an amyloid-positive status [9]. When compared with 33 controls and 18 AD dementia patients whose status was established through clinical assessment, the FreeSurfer analysis revealed a leftward asymmetry in ventral thalamic nuclei volume in the dementia group only. Stronger evidence is therefore needed to clarify thalamic changes in biologically-defined AD.

A positive amyloid PET scan is a stand-alone diagnostic tool for biologically defined AD [24]. In cognitively impaired individuals, this Core 1 biomarker usually signals that cortical AD tauopathy is also present [25]. Amyloid biomarkers become positive 15–20 years before clinical dementia [26] and are now required to initiate current disease-modifying treatments [27,28]. However, amyloid positivity alone does not fully capture the neurodegenerative process; its burden plateaus while cognitive decline progresses, and many amyloid-positive individuals remain cognitively normal for years, if not indefinitely [29,30]. Thus, complementary markers of neurodegeneration are needed to better stage disease severity and predict progression. The thalamus, vulnerable to early tau and amyloid deposition [3-7], may provide such additional information from routine MRI.

The current study examined atrophy in thalamic nuclei in 1,327 people from the ADNI dataset. Importantly, we incorporated amyloid status (positive/negative), as defined by an amyloid PET-based threshold of >24 Centiloids [31,32]. This threshold aligns with evidence of decline to dementia within five years and has been proposed as a suitable cut-off for disease-modifying intervention [27]. Combined with clinical diagnosis, this yielded six groups: amyloid-negative or amyloid-positive CN-MCI-dementia/AD. Amyloid positivity can occur in both CN subjects and non-AD dementias, reflecting either resilience or the limited specificity of amyloid PET for AD, highlighting the need for complementary biomarkers [33,34,35]. We (1) compared thalamic nuclei volumes derived from HIPS-THOMAS across these groups; (2) examined the relationship between amyloid PET load and thalamic nuclei volume; (3) assessed the diagnostic value of thalamic nuclei in a Random Forest classification; (4) constructed a Uniform Manifold Approximation and Projection (UMAP) model that integrated thalamic nuclei volumes, CSF biomarkers, cognitive scores and demographics to characterize the phenotype of individuals across the groups. We advance the premise that a subset of key thalamic nuclei can enhance current biological frameworks, improve clinical evaluation and offer new imaging biomarkers for therapeutic interventions.

## 2. Materials and methods

### 2.1. Dataset

T1w imaging data were obtained from the Alzheimer’s Disease Neuroimaging Initiative (ADNI) database (adni.loni.usc.edu; National Institute on Aging (NIH Grant U19AG024904) in compliance with the ADNI Data Use Agreement. We downloaded data from the LONI platform in July 2025. We selected isotropic (≤1mm^3^) 3D T1w MPRAGE scan acquired on a 3T scanner closest in time to the initial ADNI clinical diagnosis. Inclusion required an amyloid-PET scan within 700 days of MRI, as well as Clinical Dementia Rating (CDR), and clinical status (CN, MCI, AD) assessed on the day of MRI. Early and late MCI patients, defined only by memory impairment in a single ADNI wave, were combined as MCI. Participants were not included if they had incomplete imaging, major motion artifacts, segmentation failures, unreliable amyloid-PET score (expert rating on a standardized visual rating scale), or missing demographic/clinical information. Consequently, we restricted our analysis to ADNI2 and ADNI3 participants; other waves were precluded by insufficient group sizes and absence of key diagnostics.

### 2.2. Diagnostic Classification

Diagnostic classification for CN, MCI and AD status in ADNI participants was only based on clinical criteria per site. Clinical criteria incorporate neuropsychological testing, clinical dementia rating (CDR), and clinician judgment (see https://adni.loni.usc.edu/data-samples/adni-data/study-cohort-information/). Cognitive tests used in our analyses included Rey Auditory Verbal Learning Test (RAVLT), Trail Making Test (TMT), Boston Naming Test (BNT), clock drawing and copying, phonemic and category fluencies, global mental status (MMSE) and premorbid intellectual function (American National Adult Reading Test; ANART). CSF pTau_181_/Aβ42 ratio [36,37] were available for a subset of participants. Amyloid-PET scans in ADNI were undertaken using 18 F-Florbetaben (FBB) or 18 F-Florbetapir (FBP). Centiloid values, extracted from the ADNI database, were thresholded at > 24 to confirm an amyloid positive scan, per the Alzheimer’s Association Research Roundtable ([27, 32]; see Supplementary Table 1 for comparison with SUVR). Combined with clinical diagnosis, this Centiloid biomarker threshold was used to define six groups: amyloid-negative CN (n = 441); amyloid-positive MCI (n = 276) and amyloid-positive AD (n = 169), which together represent the biologically defined clinical AD continuum; amyloid-positive CN (n = 156), in favor of preclinical AD ; amyloid-negative MCI (n = 254) and amyloid-negative dementia (n = 31), reflecting cognitive impairment due to non-AD aetiologies. Amyloid-positive CN were deliberately not included in the “clinical AD continuum”, given ongoing debate about whether all such individuals will progress to symptomatic AD versus remain resilient [29,30]. This separation enabled us to first characterise thalamic nuclei changes in a reference group, providing a stable baseline against which to measure progressive thalamic changes in amyloid-positive MCI and AD. And second, to independently investigate whether amyloid-positive CN show heterogeneous phenotypes. This analytical choice preserves them as an independent group to investigate resilience and early neurodegeneration signatures.

**Table 1:**
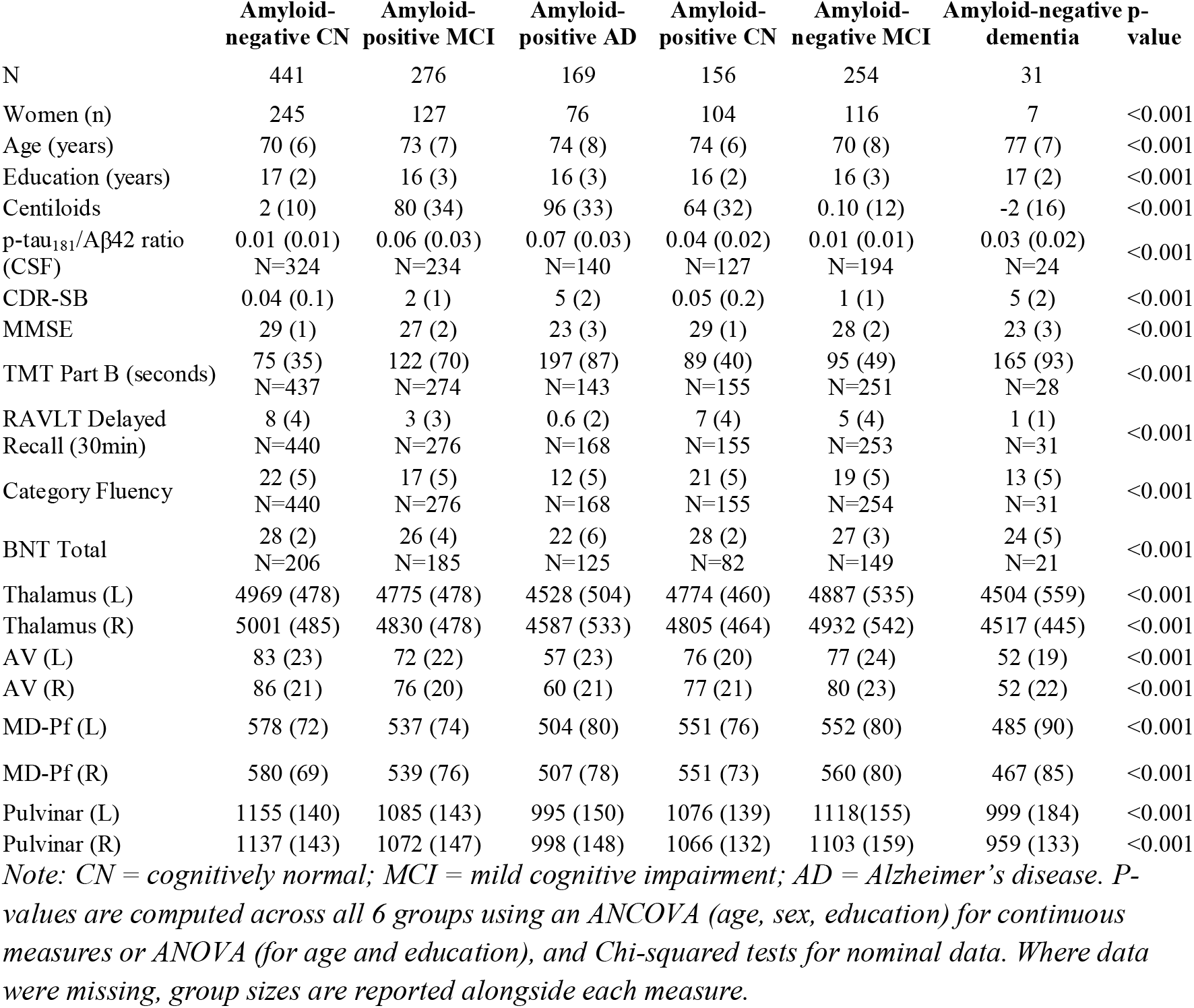
Sample characteristics.

### 2.3. HIPS-THOMAS processing

T1w MRI were preprocessed using HIPS-THOMAS [18]. This segmented 10 thalamic nuclei (anteroventral [AV], ventral anterior [VA], ventrolateral anterior [VLa], ventrolateral posterior [VLP], ventral posterior lateral [VPL], pulvinar [Pul], lateral geniculate [LGN], medial geniculate [MGN], centromedian [CM], mediodorsal including parafascicular [MD-Pf]), plus habenular [Hb], and mammillothalamic tract [MTT]. Volumes were corrected for estimated Total Intracranial Volume (eTIV) using the FreeSurfer pipeline. THOMAS includes intensity normalization and multi-atlas label fusion techniques that enhance anatomical consistency across scanners, reducing non-biological site-related variance [16,17]. NeuroCombat [38] was used to harmonize volumes across sites with age, sex, education and diagnosis group as covariates to preserve the biological variance of interest.

### 2.4. Statistical Analysis

All analyses were conducted using Python 3.12 and JASP 0.19.3.

#### (1) Comparison of thalamic volumes among groups

Each bilateral thalamic subregion was compared, first between amyloid-positive MCI and amyloid-positive AD relative to amyloid-negative CN, to assess changes along the clinical AD continuum. Second, comparisons were made between the remaining groups (amyloid-negative MCI, amyloid-negative dementia, and amyloid-positive CN) each compared to amyloid-negative CN. It aimed to assess the specificity of thalamic changes to amyloid pathology by examining non-AD cognitive impairment, and to evaluate their sensitivity in detecting preclinical AD neurodegeneration before symptom onset. Because the residuals violated ANCOVA assumptions (Levene and Shapiro–Wilk tests), we employed an Aligned Rank Transform (ART) to regress each thalamic variable on the covariates (age, sex, and years of education) and then ranked the residuals. Subsequently, Mann–Whitney U tests were used to examine these aligned ranks for the group pairwise comparisons. All p-values were corrected for multiple comparisons using the false discovery rate (FDR-corrected p < 0.05). Effect sizes for these comparisons are reported as the rank-biserial correlation.

#### (2) Association between thalamic nuclei and amyloid load

Partial Pearson correlations between Centiloids and thalamic volumes were conducted for each group using residual values after adjustment for age, sex, and education. Statistical significance was assessed with FDR correction. We also tested whether this relationship differed among amyloid-negative CN, amyloid-positive MCI and amyloid-positive AD using ANCOVA. Interaction significance was assessed via F-statistics and partial η^2^ (effect size measure), with FDR correction on p-values for multiple comparisons.

#### (3) Incremental value of thalamic nuclei for group classification

We used a Random Forest analysis (scikit-learn, Python) to assess the incremental value of thalamic nuclei volumes in the context of classification of three groups, which were: all CN (using both amyloid-negative and amyloid-positive CN), amyloid-positive MCI, and amyloid-positive AD group. This design reflects clinical reality, where amyloid status is unknown at initial evaluation for CN, while allowing identification of an AD thalamic atrophy signature in the biologically defined healthy to preclinical to clinical AD-continuum. Critically, if groups were perfectly separated by amyloid status, the model would be biased. Permutation feature importance was assessed and leveraged to sequentially add thalamic nuclei in the order of their importance to a classification model based on Centiloids and demographics (e.g., the highest-importance nucleus was added first, then the highest plus the second-highest, and so on). All model comparisons used group-stratified 5-fold cross-validation with identical splits. The final (enhanced) model was defined as the one that achieved the largest improvement in overall classification accuracy (paired t-test, FDR-corrected p-values). To characterize discriminative ability, we conducted receiver operating characteristic (ROC) analysis using a one-vs-rest approach for each class (CN, MCI, AD). For each comparison, we computed the area under the curve (AUC) for both the “Centiloid plus demographics” model versus the model that added thalamic nucleus. Significance of these ROC-AUC comparisons was assessed using a permutation-based bootstrap test (5000 permutations) with FDR-corrected p-values.

#### (4) Phenotypic profiling (UMAP)

##### Reference model construction

To establish a phenotypic landscape of the AD continuum independent of amyloid PET values, we trained a Uniform Manifold Approximation and Projection (UMAP) dimensionality reduction model using only amyloid-negative CN (n = 180), amyloid-positive MCI (n = 171), and amyloid-positive AD (n = 92), the subset with complete data on all included features. Because UMAP cannot accommodate missing values, any participant with incomplete data on the included features was excluded from this analysis. Features comprised cognitive scores (RAVLT delayed recall, TMT-B, BNT, clock drawing/copying, phonemic and category fluencies, MMSE, ANART errors), everyday function (CDR Sum of Boxes), demographics (age, education), CSF p-tau181/Aβ42 ratio, and thalamic nuclei volumes. Amyloid PET Centiloids were excluded to avoid circularity. Features were z-scored to ensure comparable contributions to Euclidean distances, and UMAP was applied to generate a 2D reference space. The resulting 2D positions for individuals reveal the spatial distributions of diagnostic groups and are also visualized as density plots using kernel density estimation that show regions of high subject concentrations. Feature importance was quantified via Pearson correlation, mutual information, and Random Forest R^2^, averaged into a combined importance score to identify the most discriminative variables.

##### Projection of other groups

Having defined this reference landscape of the clinical AD continuum, we projected three independent groups onto the same space using the identical UMAP transformation: amyloid-positive CN (preclinical AD, n = 74), amyloid-negative MCI (non-AD cognitive impairment, n = 133), and amyloid-negative dementia (n = 14) with complete data. A Euclidean nearest centroid classifier, trained on the reference groups, assigned each projected subject to a diagnostic label (CN, MCI, or AD). This approach tested whether individuals whose amyloid status and clinical diagnosis were not congruent would be reclassified based on their multimodal phenotype.

#### Analytical rationale

Longitudinal evidence indicates that not all amyloid-positive CN progress to clinical dementia [29,30]. For this reason, amyloid-positive CN were deliberately excluded from the reference continuum to separately test whether a subset of amyloid-positive CN already exhibit MCI-like phenotypes, addressing preclinical AD heterogeneity directly rather than averaging it into the continuum, conflating resilient or slowly-progressing phenotypes with early neurodegeneration. Amyloid-negative CN were retained as the reference baseline because they represent the absence of both AD pathology and cognitive impairment. Amyloid-negative MCI and dementia were projected as a separate step to assess whether non-AD cognitive impairment shares thalamic signatures with AD.

## 3. Results

Demographics and Centiloid amyloid values for the six groups are shown in Table 1. Centiloid amyloid distributions are shown in Fig. 1A, B. While 84.5% of the AD participants were amyloid positive, and 73.9% of the CN group were amyloid-negative, the proportion was almost even for the MCI clinical group with only 52.1% amyloid-positive MCI participants. The proportions are similar to a recent report for ADNI MCI and dementia participants, although we found a higher proportion of amyloid-negative CN participants compared to that study [39].

**Figure 1.**
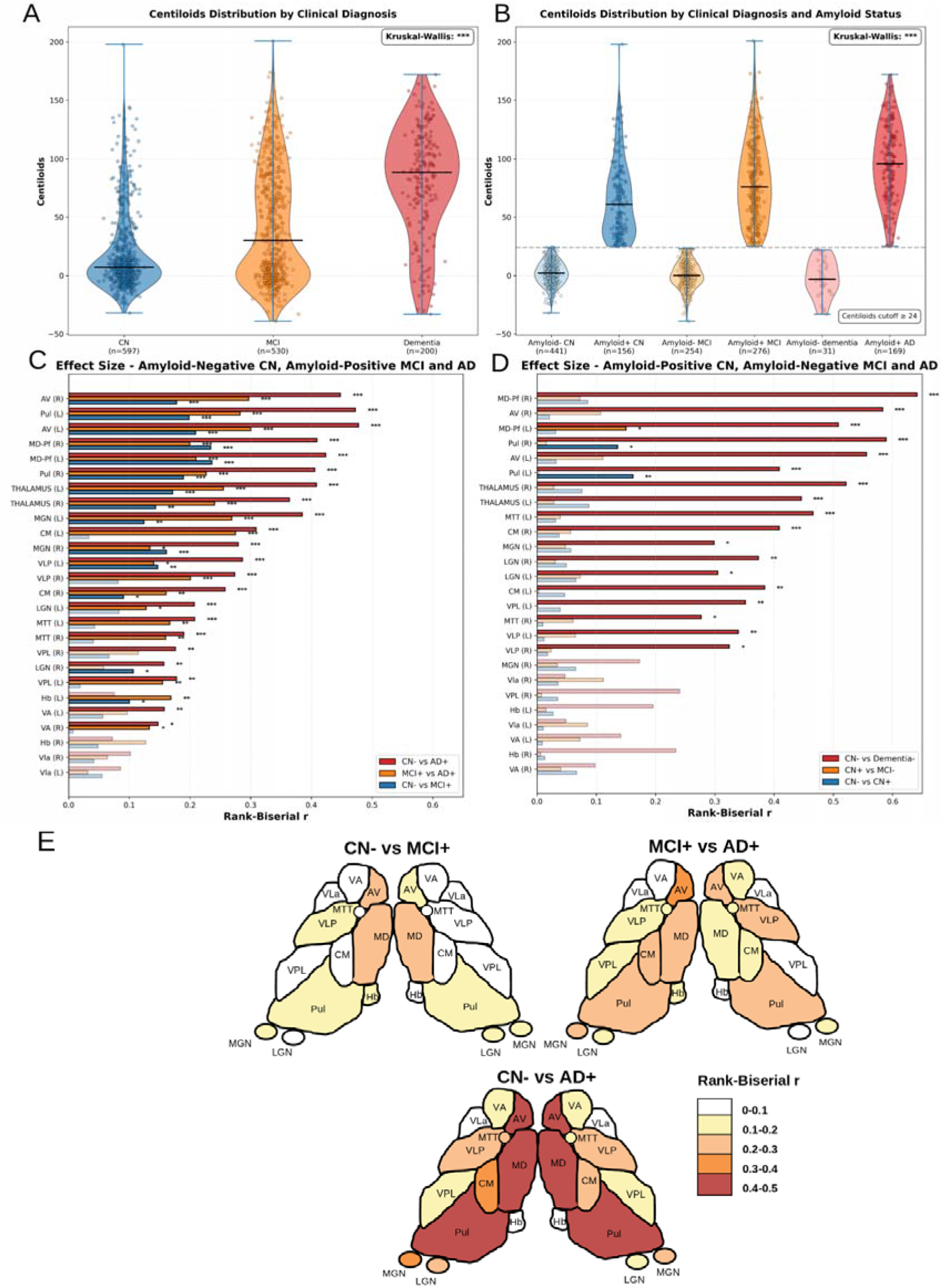
Centiloid distributions among clinical diagnostic group (CN, MCI, AD) (A) based on clinical diagnosis irrespective of amyloid status and (B) after applying the amyloid 24 Centiloid cut-off (dashed line). Amyloid-negative (Amyloid-); Amyloid-positive (Amyloid+).

### 3.1. Comparison of thalamic volumes among groups

Differences in thalamic volumes among groups are shown in Figure 1C, D, E and in Supplementary Table 2. Volumes of all thalamic nuclei were significantly different across the amyloid-negative CN, amyloid-positive MCI and amyloid-positive AD groups (FDR-corrected p < 0.05) except for the bilateral VLa nucleus and right Hb (Fig. 1C, Supp. Tab. 2). The largest effect sizes were observed for the AV, MD-Pf, and Pul nuclei bilaterally, with the left AV and MD bilaterally showing the greatest difference between amyloid-negative CN and amyloid-positive MCI. Some regions (CM, VLP, LGN, MTT, VPL, and VA) revealed small and non-significant differences between amyloid-negative CN and amyloid-positive MCI, but significant differences between both these groups and amyloid-positive AD, suggesting that they may be less involved in early progression in amyloid-positive MCI subjects (Fig. 1C). Notably, the largest difference between MCI and AD was for the AV nuclei bilaterally, suggesting its sensitivity to disease progression. An illustration summarising these results is shown in Fig. 1E.

In the amyloid-positive CN group, the pulvinar was significantly smaller compared to amyloid-negative CN (Fig. 1D), and several nuclei depicted smaller volumes (Supp. Tab. 2), suggesting possible thalamic signs from prodromal stages.

The amyloid-negative MCI and amyloid-negative dementia groups (non AD-aetiologies) also revealed thalamic nuclei differences. For amyloid-negative MCI, only the left MD-Pf had a significant smaller size, pinpointing a possible pattern in AD dementia compared to other etiologies. In contrast, the 31 amyloid-negative dementia patients showed reduced volume in several nuclei with large effect sizes for bilateral AV and MD nuclei which may reflect the impact of alternate neuropathologies (Fig. 1D).

Medians shown by black horizontal lines. (C-D) Post-hoc effect size comparison summary (Rank-Biserial r, Mann-Whitney U tests on residuals regressed for age, gender and education). FDR-corrected p-values, *<0.05, **<0.01, ***<0.001. Faded bars correspond to non-significant comparisons. (E) Summary illustration of effect size comparisons among amyloid-negative CN, and amyloid-positive MCI and AD (clinical AD continuum). The color bar represents the Rank-Biserial r from Mann-Whitney U tests on residuals regressed for age, gender, and education (as detailed in Fig. 1C).

### 3.2. Association between thalamic nuclei and amyloid load

We next examined the relationship between amyloid burden (Centiloids) and thalamic atrophy, first among the amyloid-negative CN, amyloid-positive MCI and amyloid-positive AD groups. For brevity, Figure 2 displays correlations within the left thalamus, AV, MD-Pf, and Pul nuclei which had greater between group differences in the previous analysis (Fig. 2; Supp. Fig. 1 and 2 for right and all nuclei correlations). After adjusting for age, sex, and education, several thalamic nuclei demonstrated significant interactions with variations depending on the group. The strongest and most significant association was observed in the left Pul among amyloid-positive MCI (r ≈ −0.201, p-FDR < 0.001). Only bilateral AV showed a progressively stronger negative correlation across stages (Left r: amyloid-negative CN = -0.067, amyloid-positive MCI = -0.150, amyloid-positive AD = -0.155; Right r: CN = -0.029, MCI = -0.168, AD = -0.201; Fig. 2). No significant interaction with the diagnostic group was identified.

**Figure 2.**
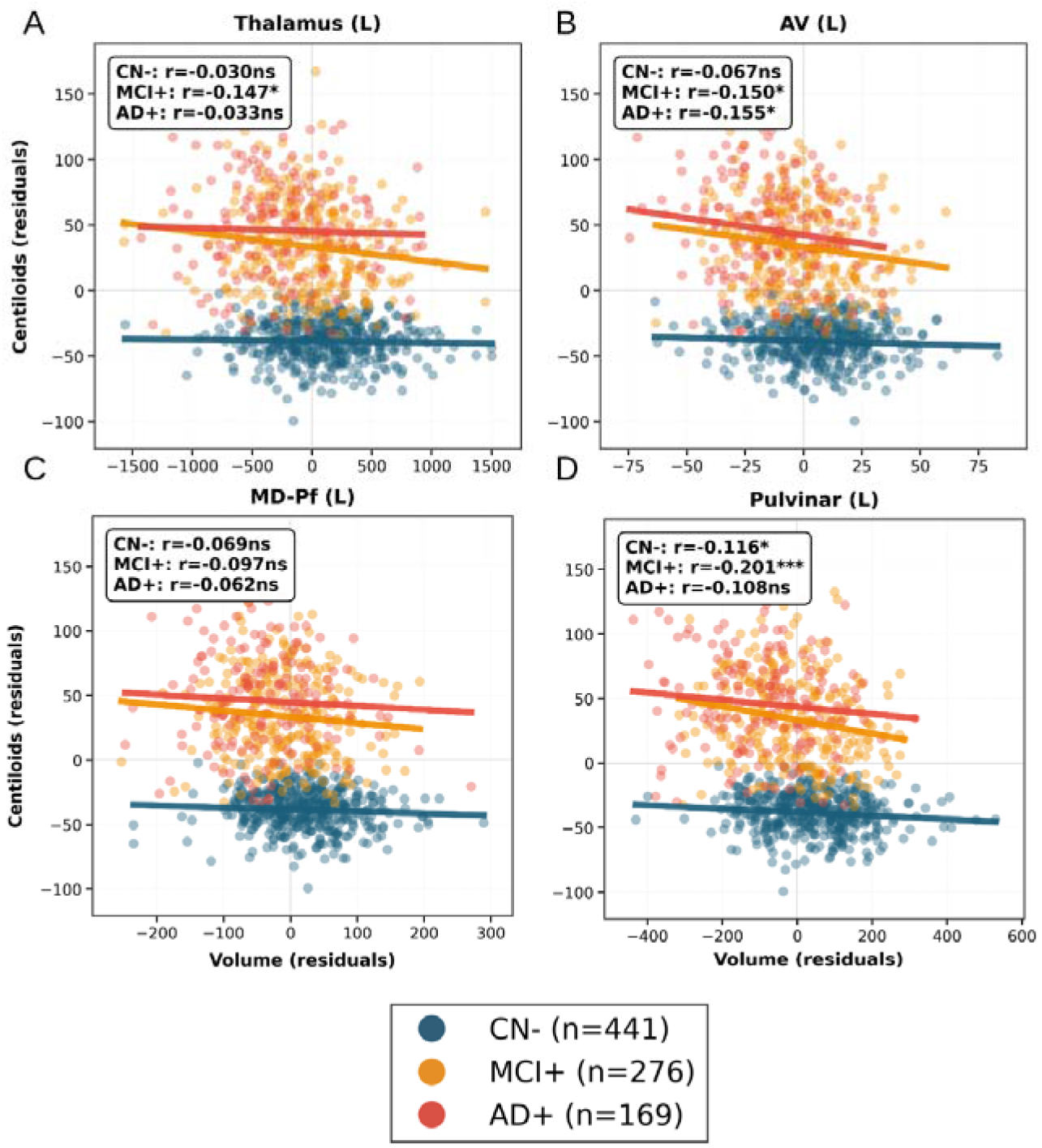
Partial correlations adjusted for age, sex, and education between Centiloid values and left (A) whole thalamus, (B) AV, (C) MD-Pf, and (D) Pulvinar volumes. Points represent residuals after covariate adjustment colored by diagnostic group: 441 amyloid-negative CN in blue, 276 amyloid-positive MCI in orange, and 169 amyloid-positive AD in red. The summary box in each panel reports the partial correlation coefficient (r), and its FDR-corrected p-value : *< 0.05, **< 0.01, ***< 0.001. ns : not significant.

We then extended this analysis to the remaining groups. Neither the amyloid-negative MCI nor the amyloid-negative dementia groups showed a significant relationship between Centiloids and any thalamic nucleus (Supp. Fig. 2).

Finally, among the amyloid-positive CN group, a significant positive correlation was found between amyloid burden and bilateral MTT (r ≈ 0.200, p < 0.05), and with the right Hb (r = 0.208, p < 0.01) (Supp. Fig. 2).

### 3.3. Incremental value of thalamic nuclei for group classification

To determine whether thalamic nuclei volumes provide additional diagnostic information beyond amyloid PET and demographics, we trained Random Forest classifiers to predict the three-class diagnosis (CN, MCI, AD) using biologically defined groups: amyloid-negative and amyloid-positive CN (n=597), amyloid-positive MCI (n=276), and amyloid-positive AD (n=169). Including both amyloid-negative and amyloid-positive CN in the control class forces the model to distinguish MCI and AD from the full spectrum of CN individuals, mirroring real-world clinical heterogeneity and reducing selection bias while amyloid-negative MCI and AD are excluded to avoid learning non AD-dementia thalamic atrophies.

Permutation importance analysis identified AV bilaterally (importance score left: 0.028, right: 0.018), left MD (0.023), left MGN (0.015) and left Pul (0.013) as the top predictive nuclei to classify the three-class diagnosis (CN, MCI, AD) (Fig. 3A). Adding left AV volume to the baseline model (Centiloids + age, sex, education) significantly improved overall classification accuracy from 69.6% to 73.0% (absolute increase of 3.4%, FDR-corrected p = 0.044, Fig. 3B). No other thalamic model yielded a statistically significant improvement after FDR correction. The improvement observed when adding AV to the model was driven primarily by enhanced classification of MCI (+2.2% AUC) and AD (+4.7% AUC) status, but not CN subjects (Fig. 4). The resulting confusion matrix of the classification is represented in Supplementary Table 3.

**Figure 3.**
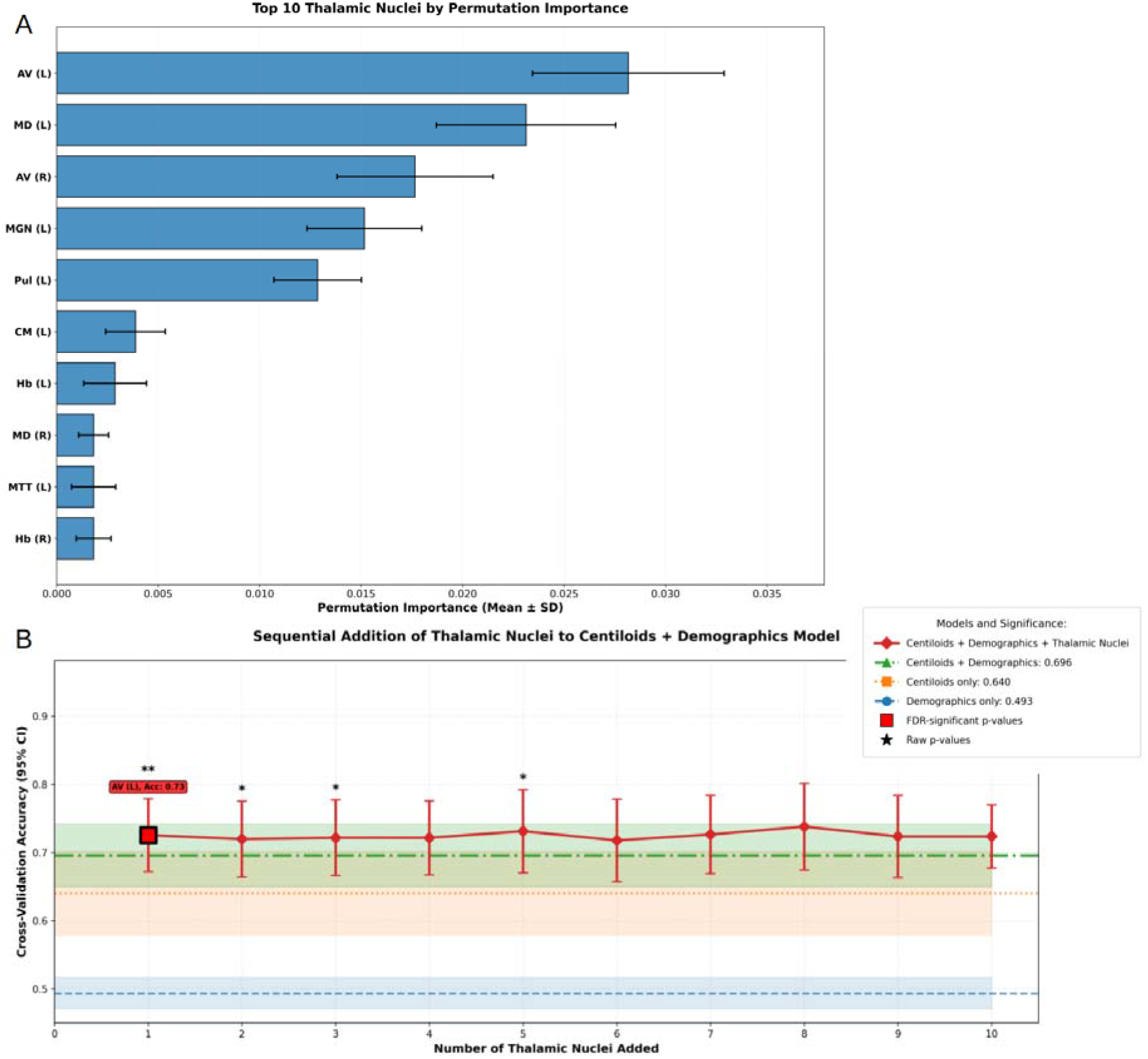
A-Mean +/-SD permutation importance on all thalamic nuclei and B-Mean cross-validation accuracy +/-95% confidence intervals by predictive models using Random Forest classifiers. Horizontal lines with shaded confidence bands represent baseline comparators: demographics only (blue dashed), Centiloids only (orange dotted), and the primary baseline of Centiloids plus demographics (green dash-dot). The solid red line illustrates the cross-validation accuracy achieved by adding thalamic nuclei, ranked by permutation importance, to the combined Centiloids and demographics model. The red square identifies the optimal model achieving FDR-corrected significant improvement while black asterisks above error bars denote raw p-value significance levels compared to the Centiloids + demographics model (*p<0.05, **p<0.01, ***p<0.001) from paired t-tests conducted on identical 5-fold stratified cross-validation splits.

**Figure 4.**
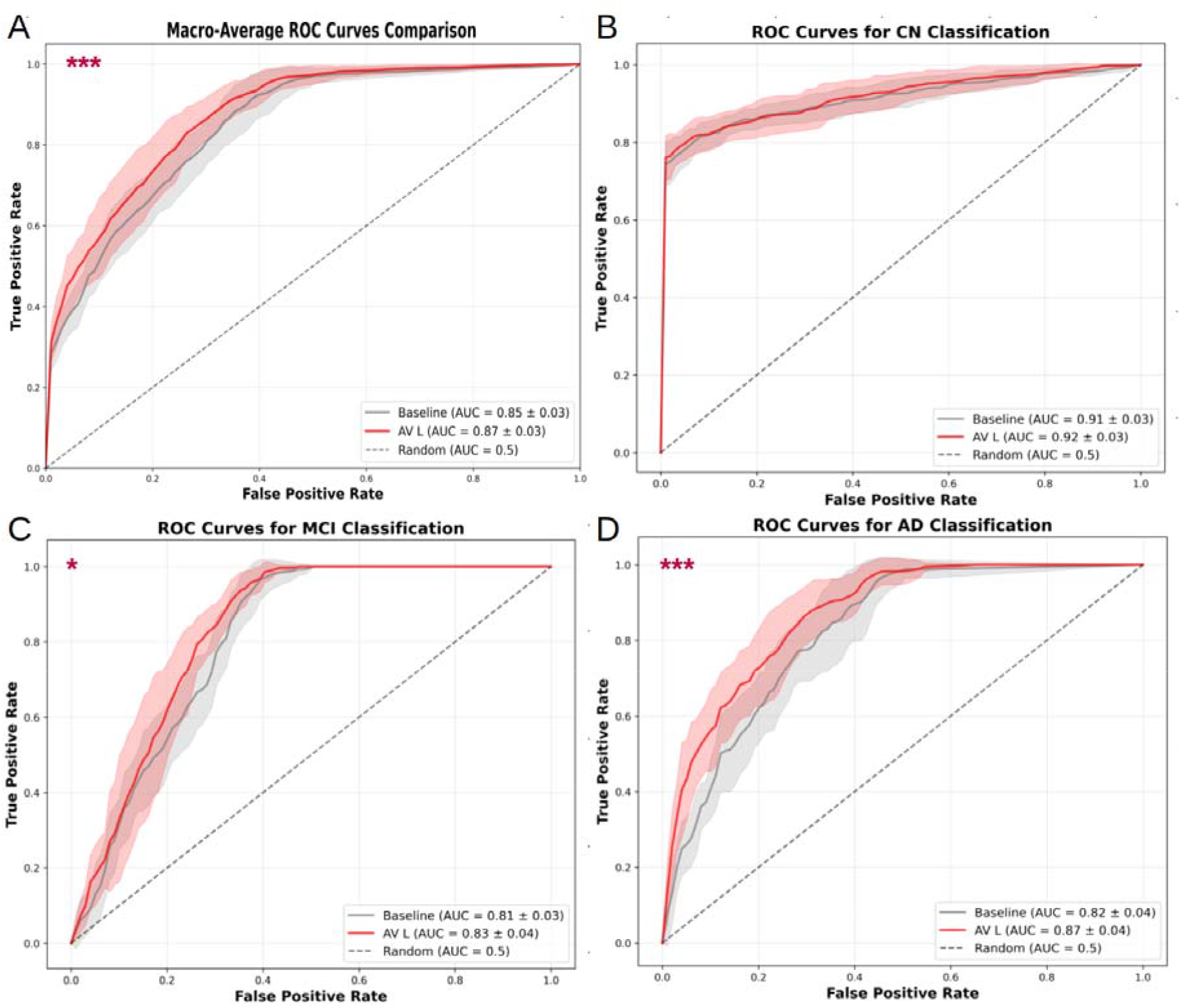
ROC Analysis (AUC ± SD across 5-fold cross-validation) comparing baseline (Demographics + Centiloids) vs enhanced models (Demographics + Centiloids + left AV) for AD stage classification using a Random Forest classifier. A. Macro-average ROC curves showing overall diagnostic performance across all classes (597 CN, 276 amyloid-positive MCI, 169 amyloid-positive AD); ROC curves for (B) CN, (C) MCI, and (D) AD classification. Gray lines represent baseline model (Centiloids + demographics); red lines represent enhanced model (Centiloids + demographics + left AV thalamic nucleus volume).

Using Centiloids and demographics alone, 80 CN subjects were classified as MCI-like and 18 as AD-like. After adding left AV volume, only 3 of these 98 misclassified CN subjects were correctly reclassified as CN. This absence of improvement in CN classification is itself informative. When the baseline model (using only Centiloids and demographics) misclassified these individuals as MCI based on their high amyloid burden, adding left AV volume confirmed rather than corrected that classification. This suggests that thalamic atrophy is already present in amyloid-positive CN, aligning with previous results (Fig. 1D ; Tab. 1).

### 3.4. Phenotypic profiling (UMAP)

In this analysis, we aimed to: 1) identify which multimodal features are most relevant for distinguishing amyloid-negative CN from the clinical AD continuum (amyloid-positive patients); and 2) clarify clinical ambiguity in cases where amyloid status and cognitive presentation are not congruent, specifically among amyloid-negative patients (cognitive impairment despite negative amyloid scans) and among amyloid-positive CN individuals (preclinical AD suggested by amyloid status despite normal cognition).

1. We constructed a phenotypic landscape using UMAP dimensionality reduction on 180 amyloid-negative CN, 171 amyloid-positive MCI and 92 amyloid-positive AD subjects, based on the subset with CSF p-tau/Aβ42 ratio, in addition to cognitive scores, demographics and thalamic volumes. Amyloid-PET Centiloid values were excluded from the model to minimise circularity, as they defined the initial groups, but also to assess the accuracy of the model in its absence. UMAP successfully separated these groups with distinct spatial distributions (Fig. 5A). The density plot revealed two strong density peaks reflecting one profile for amyloid-negative CN subjects and one for amyloid-positive AD patients. The amyloid-positive MCI participants were distributed in intermediate regions (Supp. Fig. 3). Feature importance analysis identified p-tau/Aβ42 ratio, CDR-Sum of Boxes, and MMSE as the strongest discriminators of clinical phenotype in this analysis, followed by TMT-B, RAVLT, and AV (L) volume, all with combined importance scores > 0.5 (Fig. 5B).
2. We then projected 74 amyloid-positive CN, 133 amyloid-negative MCI, and 14 amyloid-negative dementia with complete data, onto this multidimensional space. When classified on their proximity to the previous reference phenotypes (amyloid-negative CN and amyloid-positive patients), significant reclassification occurred in 27% of amyloid-positive CN subjects to MCI-like while 73% remained classified as CN, and 68% of amyloid-negative MCI were reclassified as CN. The amyloid-negative dementia patients also showed some reclassification with 7% as CN and 36% as MCI (Fig. 5C, D).

**Figure 5.**
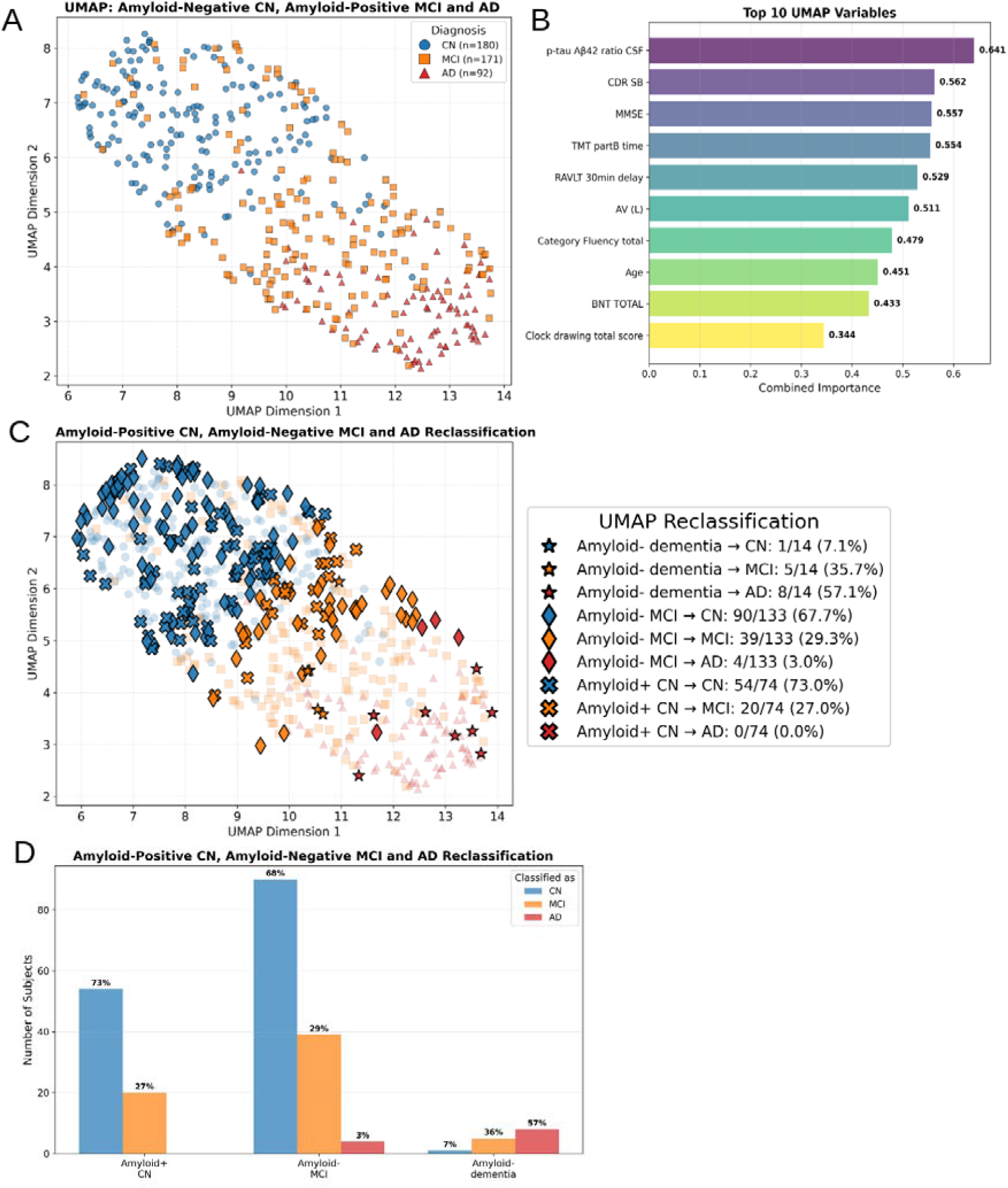
UMAP analysis. A. UMAP embedding of amyloid-negative CN (blue circles), amyloid-positive MCI (orange squares), and amyloid-positive AD (red triangles). B. Associated feature importance analysis displaying variables most strongly associated with these UMAP dimensions. The combined importance score is made of the average of mutual information, Pearson’s correlations and Random Forest regression scores. C. Reclassification outcomes of amyloid-negative patients and amyloid-positive CN when projected into UMAP space. Marker shapes indicate the initial clinical classification, while colors represent nearest centroid reclassification: blue=classified as CN, orange=classified as MCI, red=classified as AD. Background shows training on amyloid-negative CN and amyloid-positive patients at reduced opacity. D. Reclassification outcomes summary.

## 4. Discussion

This study leverages advanced thalamic segmentation (HIPS-THOMAS) to demonstrate that thalamic nuclei are central to the neurodegenerative mapping of Alzheimer’s disease (AD). By anchoring our analysis to the amyloid-PET Centiloid framework, we identified an intra-thalamic atrophy “fingerprint” that tracks with both biological pathology and clinical phenotype.

Our findings yield four primary conclusions. (1) Atrophy of nearly all thalamic nuclei escalates across the amyloid-negative CN to amyloid-positive MCI-AD spectrum, with limbic nuclei (AV, MD-Pf) and the Pul showing the largest effect sizes. Amyloid-positive CN already exhibited reduced thalamic volumes compared to amyloid-negative CN, confirming their position on the AD continuum. Amyloid-negative MCI showed no or only small effect sizes compared to amyloid-negative CN, while amyloid-negative dementia participants exhibited large volume reductions (2) Most nuclei volumes exhibited negative correlations with amyloid burden across amyloid-positive patients (3) A classification model demonstrated that the left AV nucleus adds significant value beyond amyloid PET and demographics for classifying CN, amyloid-positive MCI, and AD (4) A multivariate phenotypic model integrating cognition, CSF biomarkers, and thalamic volumes in PET-defined amyloid-negative CN, amyloid-positive MCI and amyloid-positive AD subjects successfully stratified syndromic severity. Projecting amyloid-positive CN into this space reclassified 27% as MCI-like and 68% of amyloid-negative MCI as CN-like.

### Thalamic Atrophy as a Robust and Accessible Marker of Neurodegeneration in AD

Thalamic atrophy, particularly within limbic nuclei (bilateral AV, MD-Pf) and Pul, escalates significantly across the amyloid-positive CN-MCI-AD spectrum and correlates with amyloid burden, aligning with recent studies [15,20,22,23]. These nuclei were the only ones correlating with both biomarkers (tau, p-tau, Aβ) and cognition in Bernstein et al. [15]. Unlike their study, we found AV-amyloid associations, likely due to PET-based confirmation and larger sample size.

The vulnerability of the AV, MD and Pul nuclei is pathophysiologically grounded. Post-mortem studies show early neurofibrillary tangles and amyloid deposits in anterior thalamus from early AD-dementia [3,4]. Anterior nuclei connect reciprocally to multiple regions of the episodic memory system, and unidirectionally via MTT from mammillary bodies [19,40-43]. AV and MD serve as fronto-limbic hubs supporting memory, executive function, and attention, all impaired in AD [43-46], and along with MTT injury are associated with memory deficits [19,40,41,44,47-49]. In our study, AV reduction was an early, large effect, while MTT reduction was more dementia-stage related. Pulvinar involvement likely contributes to visual, attentional, and mnemonic disturbances [50-52].

The relative sparing of motor/sensory nuclei (VA, VLa, left VPL) from amyloid correlations suggests that neurodegeneration in AD targets cognitive networks rather than the thalamus as a whole, consistent with the clinical predominance of cognitive over motor symptoms in typical AD. The positive MTT/right Hb-Centiloid correlation exclusively in amyloid-positive CN is novel, speculatively reflecting a transient cellular response (e.g., reactive gliosis) before overt atrophy.

Nevertheless, within the amyloid-positive population, thalamic volumes provide stage-specific information: reduction in amyloid-positive CN worsens progressively in amyloid-positive MCI and AD. Interpreted with amyloid status, thalamic volumetry serves as a neurodegeneration severity marker. This quick, cost-effective measure from routine T1w MRI via HIPS-THOMAS [18] complements more invasive/expensive biomarkers.

### Incremental Diagnostic Value of Thalamic Nuclei Beyond Amyloid

Neuropathology remains gold standard, but fluid/imaging biomarkers are increasingly used due to poor sensitivity of clinical diagnosis alone [24,53-55]. Yet, amyloid positivity occurs in non-AD dementias and up to 44% of CN older adults, and non-AD co-pathology is common in AD patients, highlighting the need for complementary biomarkers [33-35,39,57,58].

Our classification framework showed that adding left AV volume to demographics and Centiloids significantly improved classification of amyloid-positive MCI and AD. This 3.4% improvement in accuracy indicates that thalamic atrophy captures neurodegenerative sequelae not fully reflected by amyloid load, which often plateaus during clinical stages. AV prominence is compelling given its early AD vulnerability and reciprocal connectivity with subiculum, retrosplenial, medial temporal, and frontal cortices [3,4,58,59]. Given the modest absolute improvement, thalamic volumetry is best viewed as a sensitive, cost-effective adjunctive biomarker.

### A Multivariate Phenotype Captures Syndromic Severity Independent of Amyloid

The AT(N) research framework classifies AD using amyloid, tau, and neurodegeneration biomarkers [24,60]. Our UMAP model builds upon this by integrating cognition, CSF pathology (p-tau/Aβ42 ratio), and thalamic atrophy to classify amyloid-negative CN, and amyloid-positive MCI/AD. Beyond amyloid-PET status, this approach validated established measures (p-tau/Aβ42, CDR-SB, MMSE) and highlighted contributions of executive dysfunction (TMT-B), memory deficits (RAVLT delayed recall), and AV volume.

### Amyloid-negative Patients and Amyloid-positive CN: Clinical Ambiguity Insights Through Multimodal Phenotyping

Consistent with our analytical strategy of projecting rather than including amyloid-positive CN in the reference UMAP, the model, trained on amyloid-negative CN and amyloid-positive MCI/AD, reclassified 27% of amyloid-positive CN as MCI-like. This suggests that a subset of clinically normal individuals exhibited CSF, thalamic, and subtle cognitive features aligning more closely with MCI than CN.

This pattern was mirrored in the Random Forest classification, where including thalamic volumes did not improve classification of the CN group (amyloid-negative and amyloid-positive CN). Some amyloid-positive CN subjects were classified as “MCI-like”, which reflects biological insight rather than a model failure. These individuals already exhibit significant volume reductions, supporting the concept of preclinical AD where structural neurodegeneration precedes clinical symptoms. The subset of amyloid-positive CN who remained classified as CN may represent a resilient phenotype, potentially reflecting greater cognitive reserve or protective factors that buffer against the deleterious effects of amyloid pathology.

Recent observations suggest that higher Centiloid values increase long-term cognitive decline risk [61]. However, Dubois and colleagues [30] suggested that amyloidosis alone may be insufficient to predict progression to AD over a 30-month period. Further investigation of protective factors and co-pathologies (e.g., alpha-synuclein) is needed [56].

Similarly, over two-thirds of amyloid-negative MCI were reclassified as CN-like in the UMAP, suggesting their impairment may not reflect AD neurodegeneration. The small amyloid-negative dementia group showed mixed reclassification, highlighting differential diagnosis challenges in late-stage syndromes of dementia. These findings underscore heterogeneity within clinically defined MCI and the importance of biomarker confirmation [24,53].

Overall, this multimodal approach can refine diagnosis by identifying phenotypically resilient amyloid-positive individuals or amyloid-negative (non-AD) but impaired individuals, potentially improving patient selection for disease-modifying therapies.

### The Challenge of Specificity in Dementia Pathologies

UMAP captures a glimpse of the AD continuum, but specificity remains challenging as many features overlap with other dementias (∼30% of dementia cases are non-AD [63,64]). Tangle-only dementia [65], argyrophilic grain disease [66], Lewy body dementias/Parkinson’s disease dementia [65,67], and frontotemporal lobar degeneration [68,69] share cognitive/tau vulnerabilities, and amyloid deposition can occur in these dementias. Although amyloid contribution to cognitive deficits in those pathologies is uncertain [70]. Our finding that amyloid-negative dementia participants have large thalamic volume reductions confirms thalamic atrophy as a general neurodegeneration marker [1,2]. Future research should define disease-specific thalamic atrophy fingerprints (e.g., FTLD, Lewy body diseases), transforming thalamic volumetry into a low-cost differential diagnosis tool.

### Limitations

The analyses are cross-sectional; within-subject trajectories of thalamic atrophy relative to amyloid are needed. The small sample of amyloid-negative dementia cases prevented meaningful analysis of specific non-AD patterns. While the 3.4% improvement in overall accuracy in diagnostic classification is modest, it is statistically significant and represents information gained beyond amyloid pathology. Amyloid burden is assessed as a global load for classifications, so its analysis in specific brain regions would be insightful.

## 5. Conclusion

This study demonstrates that thalamic nuclei atrophy, especially in the anteroventral, mediodorsal, and pulvinar nuclei, escalates across the clinical AD continuum (from amyloid-positive CN to MCI to AD). Thalamic volumes provide incremental diagnostic value beyond amyloid PET and demographics, improving classification of MCI and AD stages. A multimodal phenotypic model, incorporating clinical scores, cognition, CSF biomarkers, and thalamic volumes, effectively stratifies disease severity and provides insight about amyloid-positive CN and patients with MCI or dementia that do not meet a threshold of amyloid positivity. More specifically, amyloid-negative cognitively normal individuals already exhibit detectable patterns including thalamic atrophy. These findings advocate for integrating thalamic volumetry, a cost-effective MRI-derived measure, into the AT(N) framework to enhance diagnostic precision and phenotypic understanding in AD. Prospective studies should further explore thalamic patterns to help differentiate AD from other dementias.

## 8. Data availability

Data used in this study are available from ADNI (adni.loni.usc.edu).

