## Supplementary material for "Thalamic nuclei insights into Alzheimer’s disease"

I. Supplementary materials

**Supplementary Table 1:** Cross-tabulation of amyloid status defined by ADNI protocol (SUVR-based) versus Centiloid-based classification (>24).

|  | Amyloid-  CN | Amyloid+  CN | Amyloid-  MCI | Amyloid+  MCI | Amyloid-dementia | Amyloid+  AD |
| --- | --- | --- | --- | --- | --- | --- |
| ADNI Amyloid- | 417 | 0 | 242 | 0 | 28 | 0 |
| ADNI Amyloid+ | 24 | 156 | 12 | 276 | 3 | 169 |
| **Total** | **441** | **156** | **254** | **276** | **31** | **169** |

*Note: Amyloid-negative (amyloid-) ; Amyloid-positive (amyloid+).*

**Supplementary Table 2:** Thalami and thalamic subregions bilateral volumes (mean (SD)). P-values are computed using an ANCOVA (age, sex, education) between either amyloid-negative CN, and amyloid-positive patients or amyloid-negative CN, amyloid-positive CN, and amyloid-negative patients. Pairwise effect sizes are represented in Figure 1B and 1C.

|  | **Amyloid-negative CN** | **Amyloid-positive MCI** | **Amyloid-positive AD** | **p-FDR** | **Amyloid-positive CN** | **Amyloid-negative MCI** | **Amyloid-negative dementia** | **p-FDR** |
| --- | --- | --- | --- | --- | --- | --- | --- | --- |
| **N** | 441 | 276 | 169 |  | 156 | 254 | 31 |  |
| **THAL (L)** | 4969 (477) | 4775 (478) | 4528 (504) | 2.53E-16 | 4774 (460) | 4887 (535) | 4504 (559) | 4.66E-05 |
| **THAL (R)** | 5001 (485) | 4830 (478) | 4587 (533) | 6.97E-14 | 4805 (464) | 4932 (542) | 4517 (445) | 2.48E-05 |
| **AV (L)** | 83 (23) | 72 (22) | 57 (23) | 3.98E-23 | 76 (20) | 77 (24) | 52 (19) | 3.95E-07 |
| **AV (R)** | 86 (21) | 76 (20) | 60 (21) | 5.32E-24 | 77 (21) | 80 (23) | 52 (22) | 1.09E-08 |
| **VA (L)** | 257 (34) | 252 (34) | 244 (35) | 0.006 | 249 (34) | 256 (35) | 251 (31) | 0.32 |
| **VA (R)** | 273 (38) | 270 (38) | 260 (37) | 0.011 | 268 (37) | 275 (39) | 270 (43) | 0.73 |
| **Vla (L)** | 77 (11) | 75 (11) | 75 (12) | 0.25 | 75 (10) | 76 (11) | 77 (12) | 0.30 |
| **Vla (R)** | 80 (12) | 79 (10) | 78 (12) | 0.14 | 79 (11) | 79 (11) | 81 (12) | 0.21 |
| **VLP (L)** | 771 (93) | 739 (99) | 709 (102) | 1.29E-07 | 749 (94) | 760 (106) | 707 (105) | 0.020 |
| **VLP (R)** | 800 (102) | 776 (97) | 734 (109) | 2.26E-07 | 775 (95) | 793 (110) | 737 (106) | 0.061 |
| **VPL (L)** | 288 (36) | 284 (35) | 271 (37) | 0.0013 | 276 (34) | 287 (40) | 263 (41) | 0.013 |
| **VPL (R)** | 277 (36) | 271 (32) | 259 (40) | 0.0002 | 267 (34) | 276 (40) | 262 (37) | 0.26 |
| **Pul (L)** | 1155 (140) | 1086 (143) | 995 (150) | 5.32E-24 | 1076 (139) | 1118 (155) | 999 (184) | 5.76E-07 |
| **Pul (R)** | 1137 (143) | 1072 (147) | 998 (148) | 4.85E-17 | 1066 (132) | 1103 (159) | 959 (133) | 1.63E-07 |
| **LGN (L)** | 93 (19) | 87 (18) | 82 (18) | 3.74E-05 | 86 (19) | 88 (20) | 78 (20) | 0.0073 |
| **LGN (R)** | 90 (18) | 85 (18) | 81 (19) | 0.0012 | 84 (19) | 88 (20) | 73 (19) | 0.0053 |
| **MGN (L)** | 57 (8) | 55 (8) | 51 (8) | 2.21E-13 | 54 (8) | 56 (8) | 51 (10) | 0.0042 |
| **MGN (R)** | 58 (9) | 55 (8) | 53 (8) | 1.27E-08 | 55 (8) | 57 (8) | 54 (8) | 0.06 |
| **CM (L)** | 91 (15) | 88 (17) | 79 (15) | 7.07E-09 | 85 (16) | 89 (17) | 76 (15) | 0.0087 |
| **CM (R)** | 89 (16) | 85 (17) | 79 (16) | 1.67E-06 | 84 (15) | 86 (17) | 74 (15) | 0.0013 |
| **MD-Pf (L)** | 578 (72) | 537 (74) | 504 (80) | 3.32E-18 | 551 (76) | 552 (80) | 485 (90) | 2.75E-08 |
| **MD-Pf (R)** | 580 (69) | 539 (76) | 507 (78) | 4.83E-19 | 551 (73) | 560 (80) | 467 (85) | 4.22E-11 |
| **Hb (L)** | 19 (7) | 19 (6) | 17 (8) | 0.0053 | 18 (7) | 19 (7) | 15 (7) | 0.30 |
| **Hb (R)** | 17 (7) | 18 (7) | 16 (7) | 0.14 | 16 (7) | 17 (7) | 14 (6) | 0.39 |
| **MTT (L)** | 38 (7) | 37 (7) | 34 (8) | 5.00E-05 | 36 (8) | 38 (8) | 32 (10) | 0.00025 |
| **MTT (R)** | 38 (8) | 38 (7) | 35 (9) | 0.00012 | 37 (8) | 38 (7) | 34 (11) | 0.013 |

*Note: CN = cognitively normal; MCI = mild cognitive impairment; AD = Alzheimer’s disease.*

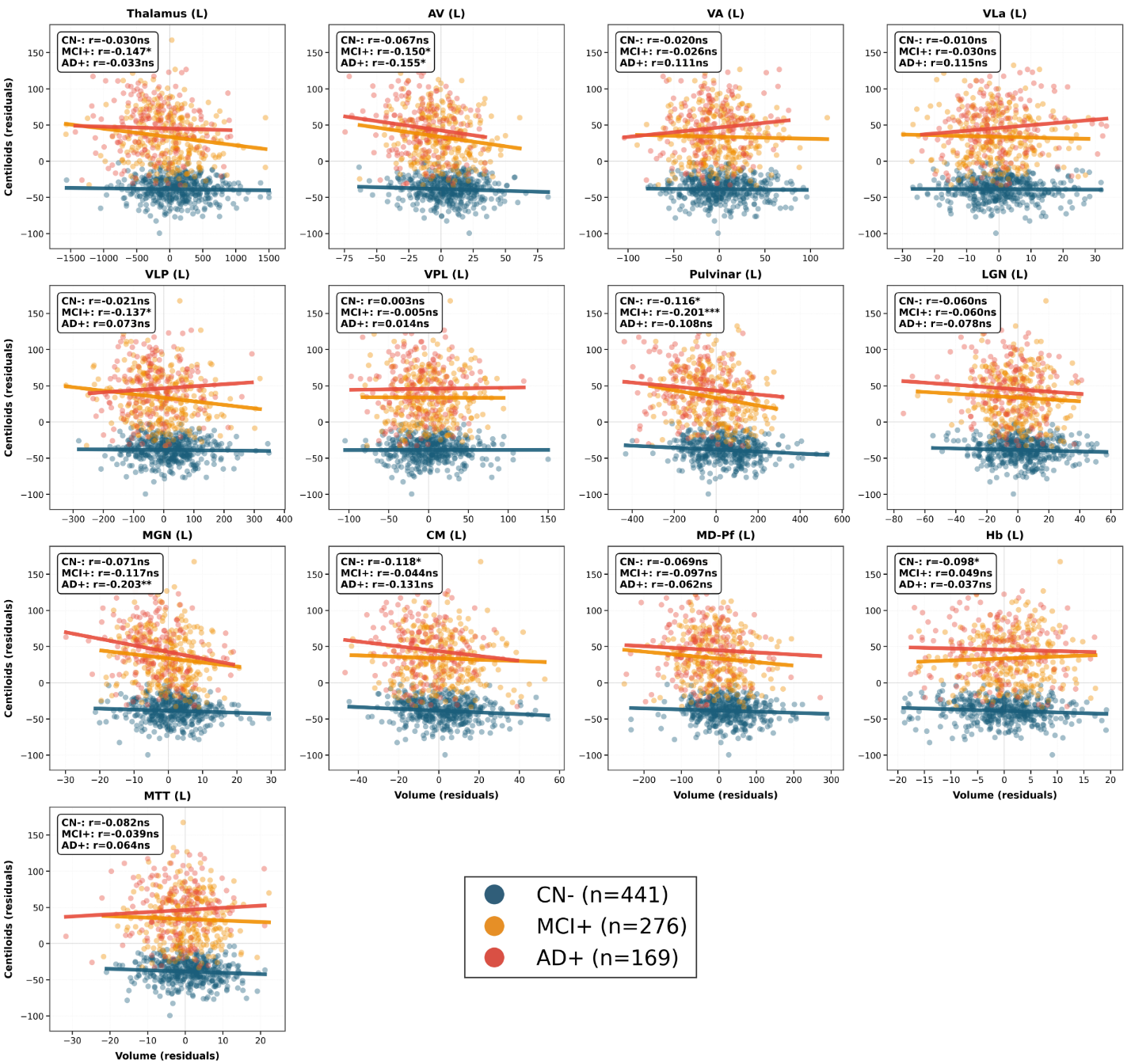

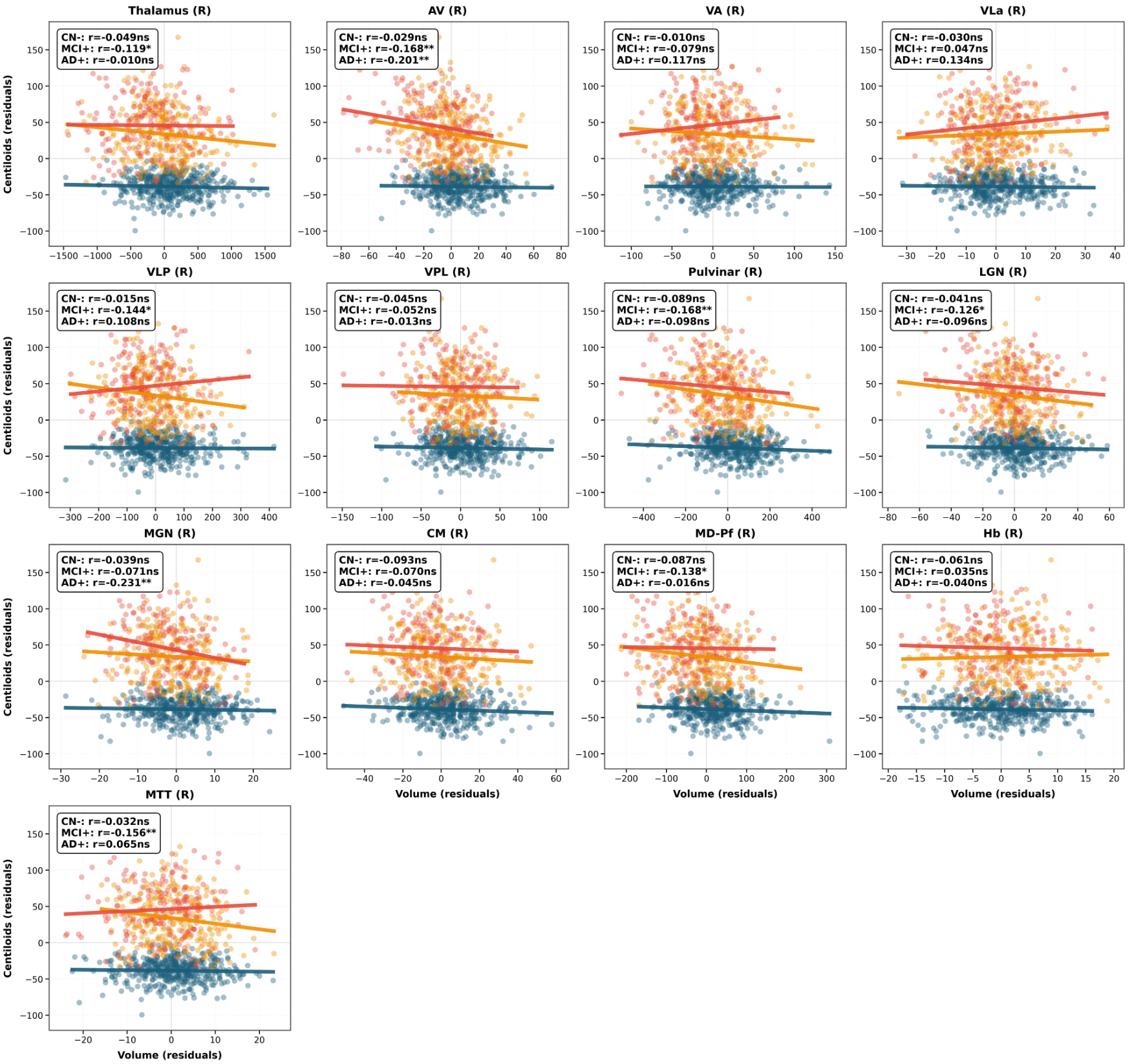

**Supplementary Figure 1:** Partial correlations between thalamic or thalamic nuclei volumes and Centiloid values, adjusted for age, sex, and education. Points represent residuals after covariate adjustment, colored by group within the clinical AD continuum (276 amyloid-positive MCI in orange, 169 amyloid-positive AD in red) compared to 441 amyloid-negative CN in blue. The summary box in each panel reports the partial correlation coefficient (r) and its FDR-corrected p-value, with significance indicated as: * < 0.05, ** < 0.01, *** < 0.001. This relationship was not significantly different depending on diagnostic group after FDR correction.

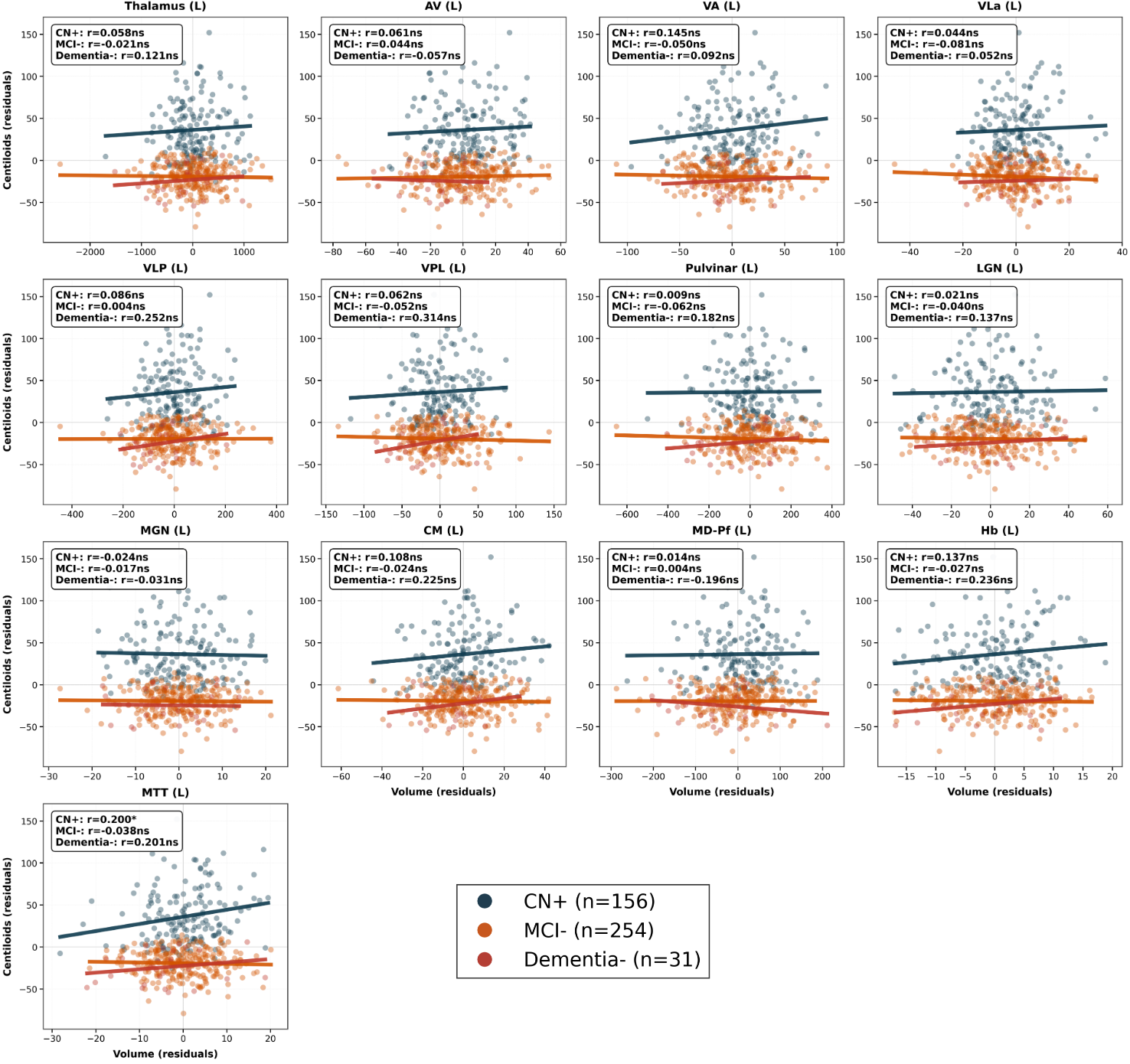

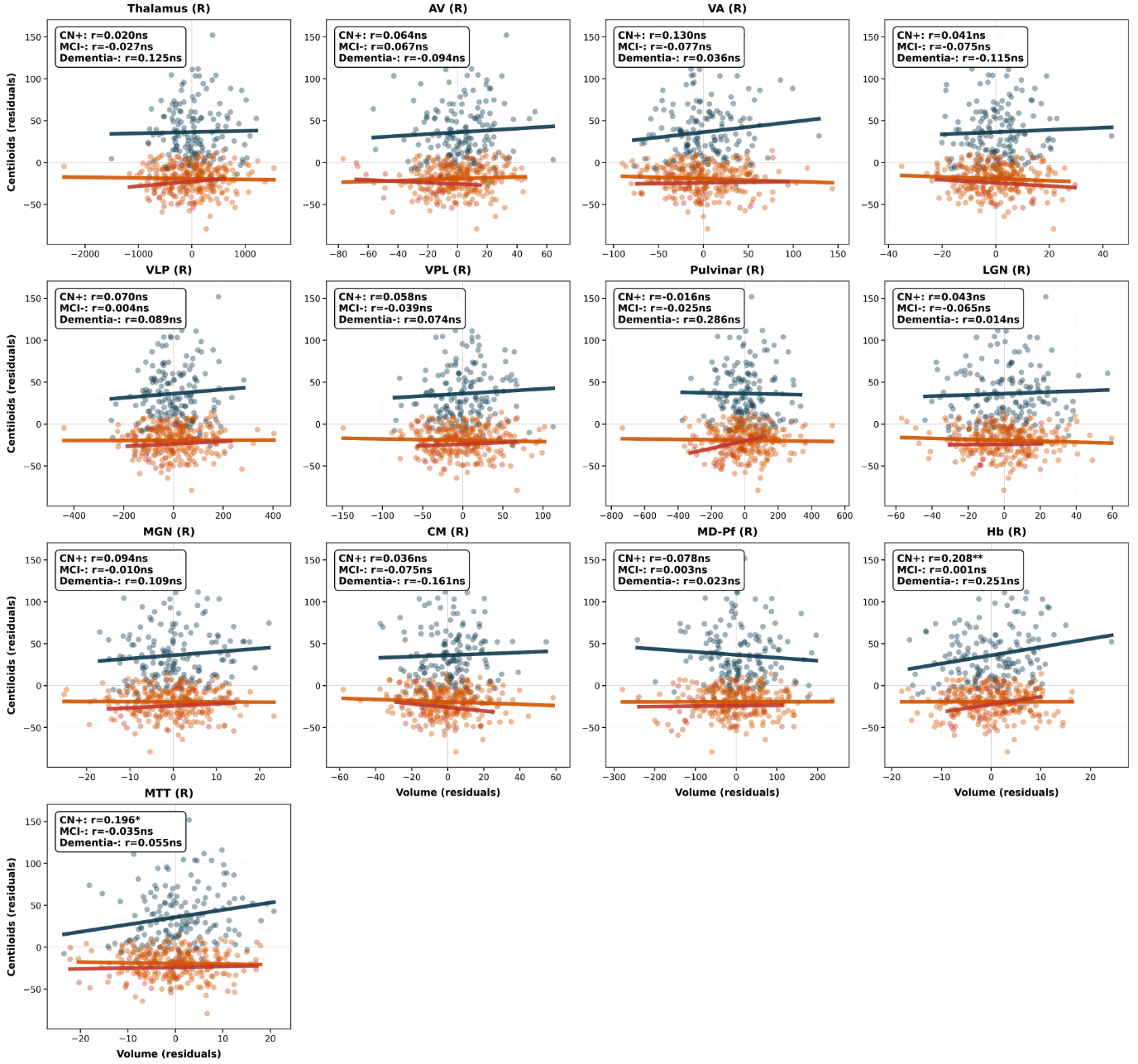

**Supplementary Figure 2:** Partial correlations between thalami or thalamic nuclei volumes and Centiloids values, adjusted for age, sex, and education. Points represent residuals after covariate adjustment colored by groups ; 156 amyloid-positive CN in blue, 254 amyloid-negative MCI in orange, and 31 amyloid-negative dementia in red. The summary box in each panel reports the partial correlation coefficient (r), and its FDR-corrected p-value : *< 0.05, **< 0.01, ***< 0.001. This relationship was not significantly different depending on diagnostic group after FDR correction.

**Supplementary Table 3**: Confusion matrix across diagnostic groups (597 amyloid-negative CN, 276 amyloid-positive MCI, 168 amyloid-positive AD) comparing baseline (Demographics + Centiloids) vs enhanced models (+ left AV volume) using a Random Forest classifier.

| Demographics + Centiloids | Pred CN | Pred MCI | Pred AD |
| --- | --- | --- | --- |
| True CN | 499 | 80 | 18 |
| True MCI | 54 | 160 | 62 |
| True AD | 19 | 84 | 66 |
| **+ AV (L)** |  |  |  |
| True CN | 502 | 79 | 16 |
| True MCI | 57 | 174 | 45 |
| True AD | 14 | 81 | 74 |

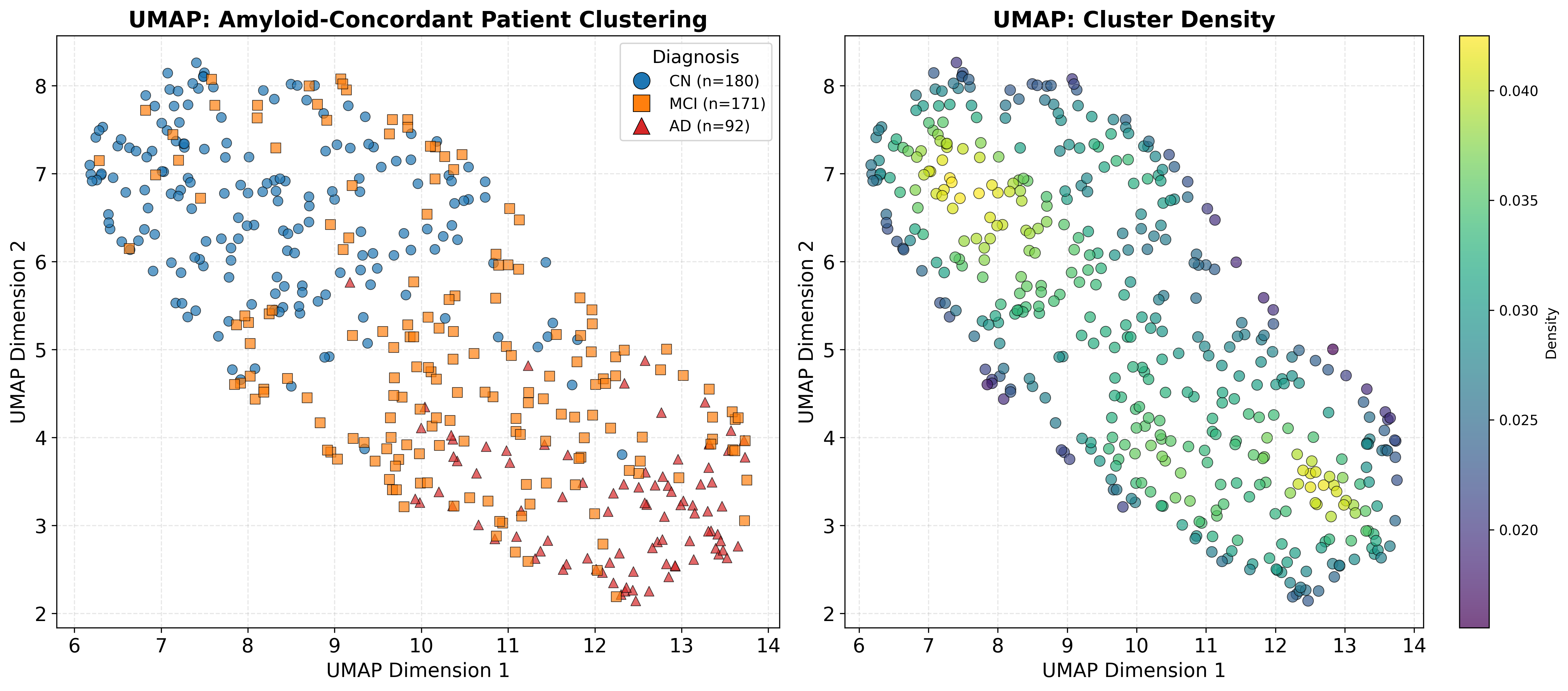

**Supplementary Figure 3:** UMAP analysis of amyloid-negative CN, amyloid-positive MCI, and amyloid-positive AD (n=443). Kernel density estimation plot highlighting two primary density peaks for CN and AD clusters.
